# Mathematical model for the distribution of DNA replication origins

**DOI:** 10.1101/2023.07.04.547751

**Authors:** Alessandro de Moura, Jens Karschau

**Affiliations:** Institute for Complex Systems and Mathematical Biology, University of Aberdeen, Aberdeen AB24 3UE, United Kingdom

## Abstract

DNA replication in yeast and in many other organisms starts from well-defined locations on the DNA known as replication origins. The spatial distribution of these origins in the genome is particularly important in ensuring that replication is completed quickly. Cells are more vulnerable to DNA damage and other forms of stress while they are replicating their genome. This raises the possibility that the spatial distribution of origins is under selection pressure. In this work we investigate the hypothesis that natural selection favours origin distributions leading to shorter replication times. Using a simple mathematical model, we show that this hypothesis leads to two main predictions about the origin distributions: that neighbouring origins that are inefficient (less likely to fire) are more likely to be close to each other than efficient origins; and that neighbouring origins with larger differences in firing times are more likely to be close to each other than origins with similar firing times. We test these predictions using next-generation sequencing data, and show that they are both supported by the data.

## I. INTRODUCTION

The life cycle of an Eukaryotic unicellular organism culminates on the S-phase, when DNA replication takes place. The cell can only proceed to cell division once all DNA in the cell has been fully replicated. This is a crucial point in the cell’s life, and there are strong selection pressures to ensure that DNA replication is as fast and error-free as possible [1, 2].

Replication starts from locations in the genome called *replication origins*[3–7]. In many organisms, the locations of replication origins are fixed, and are determined by specific DNA motifs called ARS consensus sequences (ARS stands for Autonomous Replicating Origin). This is the case with most unicellular Eukaryotes, including *S. cerevisiae*, the model organism we focus on in this paper.

A typical Eukaryotic organism has many replication origins. *S. cerevisiae*, for example, has 459 origins distributed throughout 16 chromosomes. This means that chromosomes have multiple origins, with larger chromosomes having more origins.

Before it can start replication, an origin must have been *licensed* before S-phase starts [7–12]. In *S. cerevisiae*, origin licensing is initiated by the binding of the origin recognition complex (ORC) to the origin site. In the next step, the proteins Cdc6 and Cdt1 bind to ORC, forming a complex which then loads copies of the Mcm2-Mcm7 hexamer ring (MCM) and clamps them around the DNA molecule at the origin site [13], thereby comleting the licensing process. Only licensed origins will be able to initiate replication during S-phase.

Once S-phase starts, licensing stops — depending on the organism, the licensing proteins are detached from DNA and then either degraded or inactivated [14, 15]. But some origins in any given cell will fail to license before the S-phase. Success or failure of licensing any given origin is a stochastic process: in a population of genetically identical cells, one cell may fail to license some origin, while another cell in the population may succeed in licensing that same origin [16]. This naturally leads to the definition of an origin’s *competence*, which is the probability *p*_*i*_ of origin *i* successfully licensing [16]. Alternatively, *p*_*i*_ is the fraction of cells in a population where origin *i* is licensed in a replication round.

Measuring origin competences for a whole genome is a difficult task. The duplication rate of plasmids incorporating an ARS in principle allows one to measure the competence of chosen origins [17, 18]. It would be extremely laborious, however, to use this method to obtain competences for a whole genome, and the results are not always reliable. Next-generation sequencing makes it possible to measure replication profiles with unprecedented resolution, enabling one to take genome-wide snapshots at controlled times of the state of replication of a population of cells [19]. Fitting a detailed computational model of DNA replication in yeast to this data resulted in a reliable estimation of fundamental origin parameters such as competence and mean firing time for every origin in yeast [16, 20–23]. This allows us to ask questions about how the positions of origins may be related to their firing dynamics.

The cell becomes extremely vulnerable once DNA replication starts: DNA damage caused by UV radiation and by other stresses is much more likely to kill the cell in this stage of its life cycle [24, 25], due to the induction of fork stalling and collapse. So it is in the cell’s best interests to shorten the time it spends replicating its DNA. And we know that replication origins are often created and destroyed throughout evolution [26]. This motivated us to propose, in a previous work [27], that natural selection favours origin distributions resulting in shorter replication times. We showed that this hypothesis leads to the prediction that there is a correlation between the positions of origins and their competences: neighbouring origins with low competences are expected to be located close to each other, while origins with high competence are expected to be far away from other origins. However, no genome-wide estimation of origin competences was available when that work was published; so no experimental validation was possible, and that was a mostly theoretical work.

In this paper, we use the origin parameters derived from next-generation sequencing data to quantitatively test the hypothesis proposed in [27], showing that it is supported by the data. We also extend the model presented in [27] in a major way, by incorporating the mean firing times of origins into the model. This enables us to predict a correlation between the firing times of neighbouring origins and their genomic distance. This new prediction is tested using two different next-generation sequencing data sets, and we show that it also agrees with the data, adding evidence to the idea that origin locations have been selected by evolution to favour short replication times.

## II. THE MODEL

The goal of the mathematical model we formulate here is to understand how the spatial distribution of replication origins affects the replication time; in particular, we are interested in the origin configurations that lead to the shortest replication times. We consider the simplest possible case of a chromosome with only two origins. Even though this is a very idealised model, we argue that it captures some essential aspects of how the replication time depends on the positioning of origins. The goal is to apply insights gained from this model to analyse real replication data from yeast. We generalise here the analysis done in [27], extending that model to take into account the different firing times of origins.

*S. cerevisiae* and Eukaryotes in general have linear chromosomes — as opposed to the circular chromosomes found in bacteria. So the model we develop in the following assumes linear chromosomes.

In order to simplify the model, we choose a unit of length such that the chromosome length is 1. The positions of the two origins are denoted by *x*_1_ and *x*_2_, with 0 *≤ x*_1_ *≤ x*_2_ *≤* 1. The competences of the two origins are denoted by *p*_1_ and *p*_2_. The last parameter characterising the model is the difference in the firing time between the two origins, denoted by *τ*. We assume that origin 1 is the early origin — that is, origin 1 never fires later than origin 2. This is done for mathematical convenience, and it does not affect the generality of our conclusions in any way. The replication forks are assumed to travel with constant velocity *v*, which is consistent with recent fork velocity profiles [28].

We choose *t* = 0 as the moment when origin 2 fires. Assuming both origins fire, the order of events is therefore as follows: at time *t* = *− τ*, origin one fires; at time *t* = 0, origin 2 fires; and then, the whole chromosome is replicated after all forks have either collided with other The above argument assumed both origins successfully forks or reached one of the ends of the chromosome. Notice that if a fork reaches an origin before it replicates, the fork goes through the origin and continues in its way; the origin is said to have been “passively replicated” by the fork.

This process is illustrated in Fig 1(a). We assume that *τ ≥* 0 ; that is, origin 1 either fires earlier than origin 2, or fires at the same time as origin 2. *τ* is therefore the difference in firing times of the two origins. For the purposes of this simple model, we assume that the firing times of both origins are constant, and subject to no stochastic variation. In reality, origin firing times are stochastic [16, 21, 22, 29, 30], but we ignore this at first for the sake of simplicity. We do investigate the case of stochastic firing later on in this paper.

**FIG. 1.**
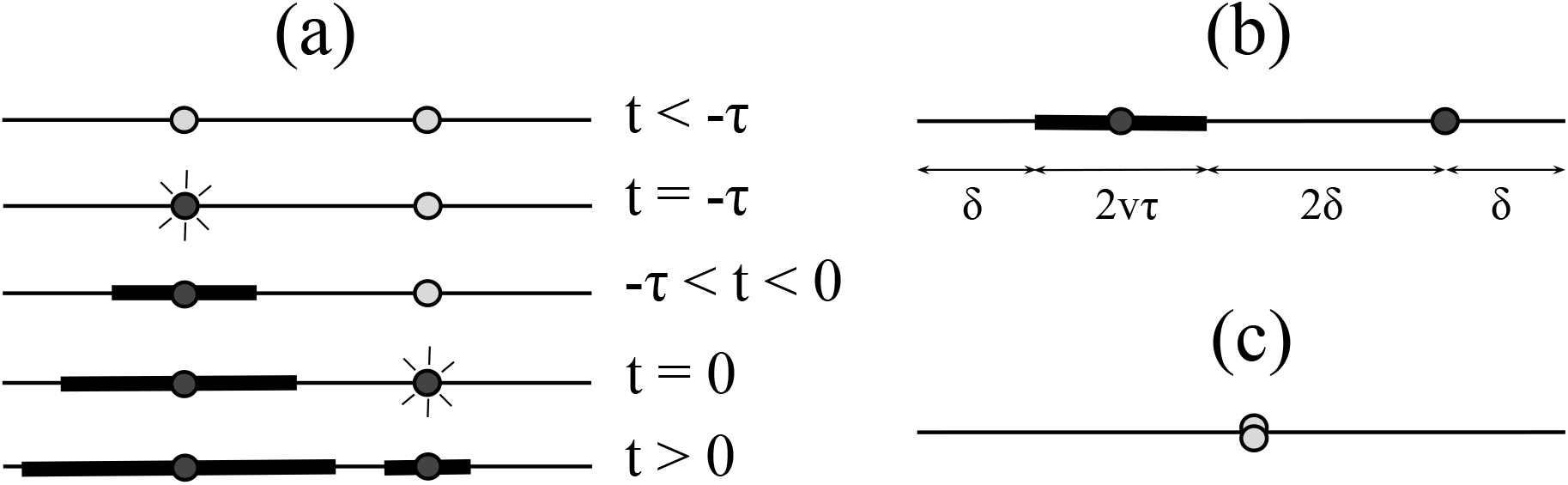
Depiction of idealised two-origin chromosome. **(a)** Sequence of origin firings in the two-origin chromosome of the model. The horizontal lines represent the chromosome, and the thick lines are the regions that have been replicated. Time progresses from top to bottom. **(b)** Minimum replication time chromosome, assuming both origins fire, depicted at *t* = 0. **(c)** Clustered origin configuration, with both origins located in the centre.

### A. Configurations of minimum replication time

We now proceed to determine where the origins should be placed so that the replication time is shortest. Let us first assume that both origins successfully fire (that is, both have been licensed). Let us focus on the moment when origin 2 (the later origin) fires, at a time *τ* after origin 1 has fired. At this time, a portion of the chromosome has already been replicated by the forks created at origin 1, as indicated by the thicker lines in Fig. 1. The question is then how to place the origins so that the remaining unreplicated portion of the chromosome is replicated as fast as possible.

The first step is to notice that after origin 2 fires, there are four forks travelling on the chromosome, and so the rate with which replication is proceeding is 4*v* (in replicated length per second). This will remain the replication rate as long as there are four forks on the chromosome. If, for example, origin 2 is very close to the right edge of the chromosome, its right-propagating fork will hit the edge and disappear soon after origin 2 fires. In that case, only three forks will remain, and replication will take place at a lower rate than the maximum possible rate, because of the absence of one fork.

The conclusion is that in order to get the shortest possible replication time, all four forks must coexist until the end of replication, so that replication remains at its fastest rate until it finishes. In other words, in the optimal configuration all forks must terminate simultaneously. This implies that the origin positions must be such that each fork will travel one-fourth of the remaining chromosomal length yet to be replicated. This reasoning leads to the origin placement shown in Fig. 1(b), where *δ* is defined as one-quarter of the unreplicated length of chromosome at the time origin 2 fires. Notice that in this configuration, the unreplicated region between the two origins is twice as large as the regions next to the edges. The reason is that the central region will be replicated by two forks, whereas the left and right regions are replicated by only one fork each.

The above argument assumed both origins successfully fired. In reality, however, each origin has a probability of firing, given by its competence. The natural quantity characterising replication time is therefore the mean replication time, which we denote by 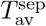. It takes into account the probabilities of all the possible combinations of origins firing and failing (we will discuss later how we deal with the awkward case of none of the origins firing.)

If both origins always fire, Fig. 1(b) is the configuration with the least average replication time. If the origins have a substantial probability of failing, however, the configuration shown in Fig. 1(b) will no longer be the one with the least replication time. The reason is simple: if one of the origins fails to fire, the origin that does fire is closer to one end of the chromosome than the other, and so one fork will terminate before the other. This violates the condition for minimal replication time, as explained above. If we knew ahead of time that only one origin would fire, the origin placement leading to the shortest replication time would be the clustered configuration depicted in Fig. 1(c), with both origins located at the centre of the chromosome. This configuration ensures that both forks created by the one firing origin terminate simultaneously. We thus expect to see a transition in the optimal origin configuration (that is, the one with the least replication time) from Fig. 1(b) for high competences to Fig. 1(c) for low competences.

Consider now the effect that the difference in firing times has on the optimal configuration, assuming for the purposes of this discussion that the origins have high competences. If the two origins fire at almost the same time, then from the arguments presented above the configuration in Fig. 1(b) is the one with minimal replication time. If, however, one of the origins is much earlier than the other, so that the later origin is passively replicated by a fork from the earlier origin before it has a chance of firing, this changes. In that case, replication will mostly be done by the earlier origin, and for the purposes of computing the replication time, it is as if the later origin did not exist. By this argument, the case of a large firing time difference *τ* is equivalent to the case of one single firing origin in the previous paragraph, and we expect the clustered configuration of Fig. 1(c) to be the one with the shortest replication time. We therefore expect to see a transition in the shortest-time configuration, from isolated to clustered origins, as *τ* increases.

We now proceed to investigate these transitions in detail through a simple mathematical model.

### B. The case of separated origins

First, we assume that both origins have competence 1, and never fail to fire. As explained above, in this case the configuration of minimum replication time is described by Fig. 1b. By time *t* = 0, each fork that started at origin 1 (the earlier-firing origin) will have travelled for time *τ* with velocity *v*. Therefore, the length of the replicated region at time *t* = 0 is 2*vτ* ; see Fig. 1(b). So at *t* = 0, the total length of the unreplicated part of the chromosome is 1 *−* 2*vτ*. Therefore, the quantity *δ* in Fig. 1(b) is given by

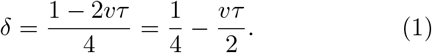

The locations of the two origins in Fig. 1(b) are thus

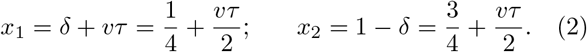

If the origins fire simultaneously, the time difference *τ* is zero. In this particular case, we find from Eq. (2) that the locations of the origins which minimise replication time are *x*_1_ = 1*/*4 and *x*_2_ = 3*/*4, where they are positioned symmetrically around the centre of the chromosome. We thus recover the result presented in [27], which assumes simultaneous firing of origins. The novel feature of our more general model is in the terms containing *τ* in Eq (2). They show that the effect of non-simultaneous firing is to break this symmetry and displace both origins so that the earlier origin (origin 1) moves closer to the centre of the chromosome, while preserving its distance from the later origin (origin 2).

The chromosome shown in Fig. 1(b) will finish replicating when each fork traverses a distance of *δ*. Since our “clock” starts when origin 2 fires, this replication time is

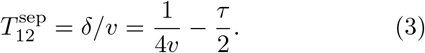

The “12” subscript indicates that this is the replication time assuming both origins fire, and “sep” indicates that we are considering the case where the origins are spatially separated.

Now let us drop the assumption that origins have competence 1. If the origin locations are still as shown in Fig. 1(b), but now origin 2 fails to fire, the new replication time is

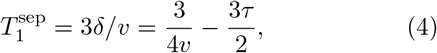

since the entire region to the right of origin 1 is replicated by the right-propagating fork originated at origin 1, which must traverse a distance of 3*δ* to reach the end of the chromosome. Similarly, if origin 1 fails, the region to the left of origin 2 is now entirely replicated by the fork originating there, and the corresponding replication time is

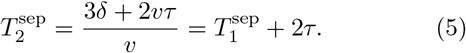

Using the definitions of the origins’ competences, *p*_1_ and *p*_2_, we can write the expression for the average replication time 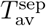

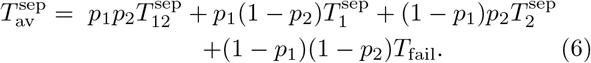

The first three terms on the right-hand side are the contributions to 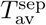 from the three scenarios described above: both origins firing; only origin 1 firing; and only origin 2 firing. The fourth term takes care of the case when both origins fail. That case would correspond to an undefined (or infinite) replication time, and it would make the definition of 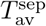 awkward. So in order to make 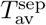 well-defined, we postulate that the case where both origins fail corresponds to some constant replication time, which we denote by *T*_fail_. This is justifiable because this is not meant to be a fully realistic model of DNA replication in yeast; it is rather meant to capture fundamental aspects of the replication dynamics. In real organisms with many origins per chromosome, the region near two unlicensed origins will eventually be replicated by forks originated from other origins. Hence, it makes sense to use a finite value for *T*_fail_. The precise value is not needed for any of the predictions we make from the model: we shall see shortly that *T*_*fail*_ cancels out in all the relevant equation

Substituting Eqs(3), (4)and (5) into Eq(6) we get

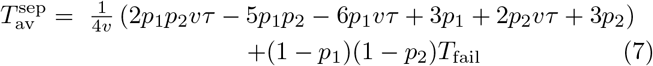

### C. The case of clustered origins

As argued above, it is reasonable to expect that if the origins have low enough competences, the configuration with both origins clustered at the centre of the chromosome (see Fig. 1c) will have the shortest replication time. We now proceed to find expressions for the replication time of the clustered case, mirroring what we did above.

Since both origins now occupy the same position, if the earlier origin (i.e. origin 1) fires, that corresponds to both 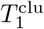 and 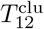 — since origin 2 is then passively 1 12 replicated by the forks created by the earlier origin. Both forks traverse half the length of the chromosome, and therefore they take the time 1*/*2*v* to finish. But they started at time *t* = *−τ*, so we have

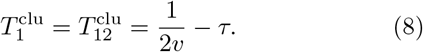

If the earlier origin fails and the later origin fires, we have instead

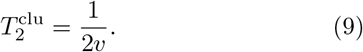

This leads to the expression for the average replication time for the clustered configuration:

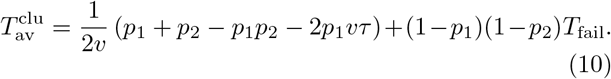

### D. Transitions in the shortest-time origin configuration

As the competences of origins decrease, we expect to see a transition of the configuration with the shortest replication time, from one with separate origins to one where the origins are clustered. That is, for high *p*_1_ and *p*_2_, we expect 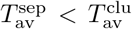, whereas for low *p*_1_ and *p*_2_, we expect 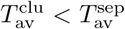. The transition point is therefore determined by 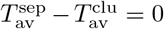. Substituting Eqs (7) and, and solving for *p*_2_, we find

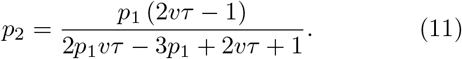

Notice that the terms involving *T*_fail_ drop out; as promised, *T*_fail_ has no effect on the conclusions that follow.

An interesting case is when *p*_1_ = *p*_2_ *≡ p*_*c*_. Then Eq. (11) becomes an equation for *p*_*c*_ and *τ*. Solving for *p*_*c*_, we get

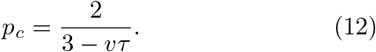

For *p < p*_*c*_, the clustered configuration is the one with the shortest replication time, whereas for *p > p*_*c*_ separate origins replicate the chromosome faster. Eq. (12) shows that *p*_*c*_ increases as *τ* increases, and hence greater differences in the firing times of neighbouring origins favour the clustering of origins. Eq. (12) can be visualised as the phase diagram shown in Fig. 2, depicting the division of the (*τ, p*) parameter space into a region where clustered origins or isolated origins yield minimal replication times. For the particular case *τ* = 0, we recover the result *p*_*c*_ = 2*/*3 found in [27].

**FIG. 2.**
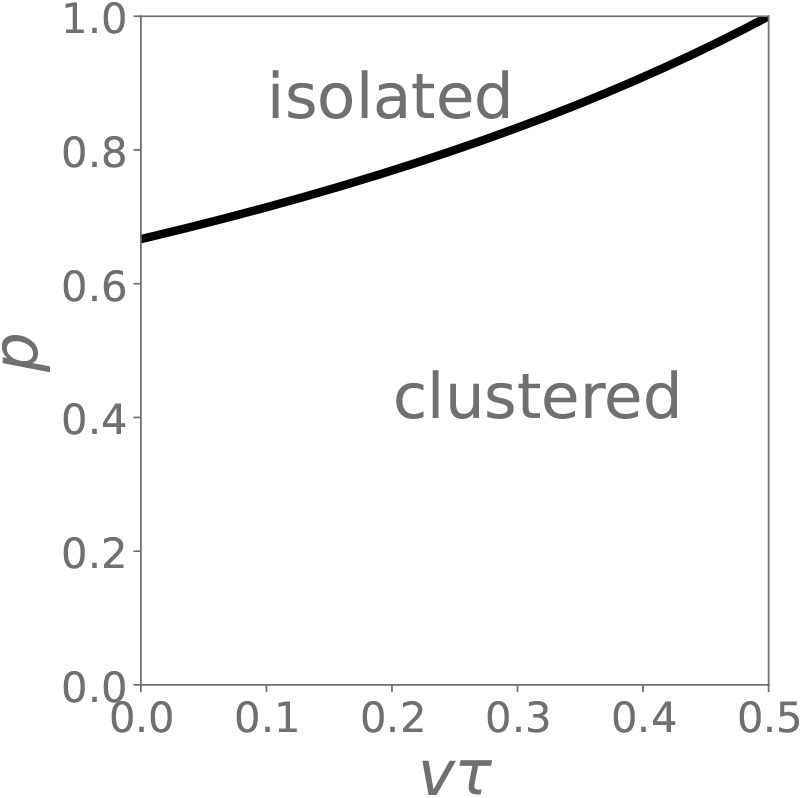
Phase diagram for the optimal time configuration of two-origin chromosomes. The horizontal axis is the difference in replication times between their origins, and the vertical axis is their competences. Above the black line, isolated origins yield minimal replication time; below it, a clustered configuration with origins very close together is optimal.

An important point is that the transition between the clustered and separated states is always abrupt — the origins do not gradually approach each other as *p* decreases. Instead, in the minimum replication time configuration, they are either in the configuration shown in Fig. 1(b) or in the clustered state shown in Fig. 1(c); there is no intermediate state. The reason for this is explained in [27]. We confirm this through the numerical simulations described in the following.

### E. Numerical simulations

In order to confirm the transitions between the clustered and isolated configurations predicted above, we ran extensive numerical simulations of the replication dynamics.

For a given origin configuration (*x*_1_, *x*_2_), the program generates a population of “virtual chromosomes”. For each member of the population, a random number *r* between 0 and 1 is generated for each origin, and then compared to that origin’s competence *p*_*i*_; if *r < p*_*i*_, the origin is considered to have successfully licensed. Then based on which origins fired, and the firing times and positions of the origins, the replication time is computed. By averaging the firing times over the virtual population, the average replication time for the given origin positions, competences and firing times are found. For given competences *p*_1_ and *p*_2_, the optimal locations of the origins are found by computing the average replication time on a 200 *×* 200 grid on the (*x*_1_, *x*_2_) space, and choosing the point on the grid yielding the minimum replication time. For convenience, we use the constraint *x*_1_ *≤ x*_2_; this takes advantage of the symmetry of the system to halve the number of points on the grid we need to consider. The size of the virtual population was 10000. The fork velocity is fixed throughout at *v* = 1, and for simplicity the origins were assumed to have the same potential *p* = *p*_1_ = *p*_2_.

We ran simulations for both constant and stochastic firing times. In the simulations using stochastic firing times, after determining whether each origin has been licensed, its firing time is chosen from a normal distribution. The mean of the normal distribution plays the role of the origin’s replication time in this version of the model. The standard deviation of the firing time distributions is constant and equal to 0.1.

The results for constant firing times are shown in Figs. 3a and 3b; the results with stochastic firing times are displayed in Figs. 3c and 3d. It is clear that transitions from clustered to isolated configurations take place as *p* increases (Figs. 3a and 3c), and as *τ* decreases (Figs. 3b and 3d), as predicted by our theory. For constant firing times, we confirmed that the values of the parameters where the transition takes place match the values predicted by the theory. And although we do not predict analytically the transition point for the stochastic case, the transition still happens, in the direction predicted by the model. This shows that the clustered-isolated transition is robust, and is present regardless of the details of the model. This gives us some confidence that our model captures fundamental aspects of the replication dynamics.

**FIG. 3.**
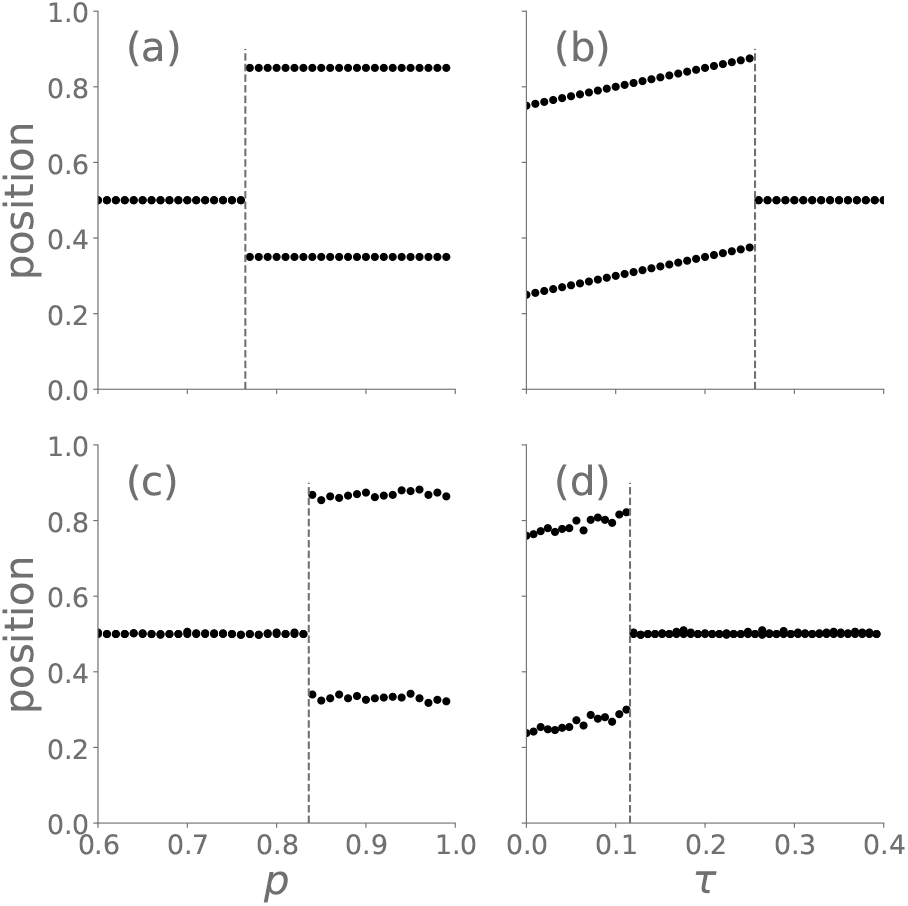
Simulations of the optimal configurations of origin positions in an idealised two-origin chromosome. *v* = 1 throughout, and the two origins have the same potential, *p* = *p*_1_ = *p*_2_. **(a)** Optimal positions versus the potential *p*, for a fixed firing time difference *τ* = 0.2, and constant firing times. **(b)** Optimal positions versus the firing time difference *τ*, for a fixed potential *p* = 0.8, and constant firing times. **(c)** Same as (a), but with stochastic firing times, with a standard deviation of 0.1. **(d)** Same as (b), but with stochastic firing times, with a standard deviation of 0.1.

### F. Predictions inferred from the model

When using the results of the analysis of the replication dynamics of the idealised two-origin chromosome to make predictions about real organisms, details such as the values of the parameters predicted by the model for the transition from the clustering to the isolated configuration should be taken with a grain of salt; this is a very idealised model of DNA replication, after all. However, its overall predictions are robust and do not depend strongly on the details of the model. In the Introduction we argue that, all other factors being equal, a shorter S-phase is advantageous, and would be favoured by natural selection. This implies that we should expect to see a significant statistical trend in favour of origin configurations leading to shorter replication times. This leads us to two main predictions:

a. The typical distance between neighbouring origins with high competence should be greater than the distance between low-competence origins;
b. The typical distance between neighbouring origins with large firing time differences should be shorter than the distance between origins with small firing time differences.

Notice that we do not expect *all* pairs of low-competence origins to be very close to each other: many factors other than the replication time are expected to affect the evolution of origin placement, and it would be unrealistic to expect the correlations found in real data to match perfectly those predicted by our model. However, we do expect to see statistically significant trends in the origin distribution in the direction indicated by these two predictions.

## III. COMPARING MODEL PREDICTIONS WITH EXPERIMENTAL DATA

We used two genome-wide origin data sets in order to compare the two main predictions of the model to available data. The first is reported in [20], and consists of the positions of all origins in *S. cerevisiae*, along with a number of origin parameters obtained by fitting next-generation sequencing replication profiles [19] to a whole-genome mathematical model of replication. For our purposes, only the location, competence and mean replication times of each origin are relevant. The second data set is taken from [23]. This data does not include competence data, but it does include firing time data. It also differs from the data reported in [20] in that it uses a slightly different set of origins for its fitting.

### Neighbouring origins with low competence tend to cluster together

In Fig. 4a, we plot the distances between pairs of neighbouring origins versus the average of the competence of the two origins in each pair. Each point in Fig. 4a represents a pair of neighbouring origins, with origin data taken from all chromosomes; in total, 443 pairs are plotted, resulting from 459 origins distributed in the 16 chromosomes of *S. cerevisiae*. One prediction from our model is that there should be a bias in the distribution of origin placement favouring short distances for pairs of low-competence origins, and larger distances for pairs with high-competence origins. This correlation is indeed seen in the figure: there is a clear trend towards a greater horizontal spread of points for greater competences, establishing that the average distance between origins grows with the average competence.

**FIG. 4.**
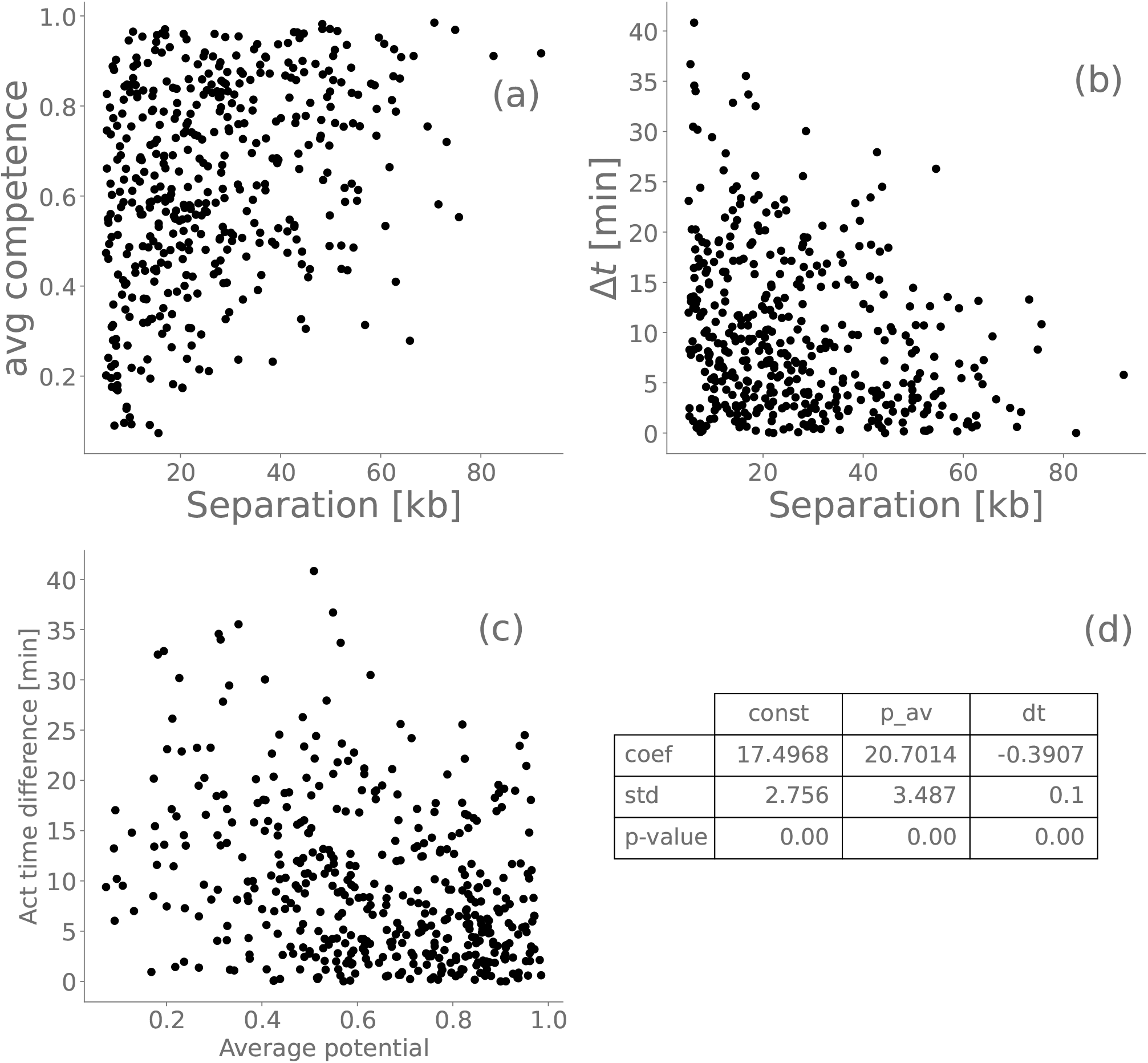
In (a), (b) and (c), each point represents one pair of neighbouring origins in *S. cerevisiae*. In (a), the horizontal axis is the distance between the origins in each pair, and the vertical axis is the average of their competence. In (b), tho horizontal axis is as in (a), and the vertical axis is the difference in average activation time between the two origins. (c) shows the mean competences versus activation time differences for each pair. Finally, (d) shows the results of multi-variable linear regression for the inter-origin distances, using the activation time differences (**dt** on the table) and mean competences (**p av**) as predictor variables. For each predictor variable, the value of the fitted coefficient, its standard deviation, and the corresponding p-value are listed. The data is taken from [20].

Another way to see this correlation is by classifying the nearest-neighbour pairs into three categories: the *high-high* pairs consist of pairs where both origins have competences greater than some threshold value *p*_0_; the *low-low* pairs are those where both origins have competences below *p*_0_; and the *high-low* pairs are the ones with a low- and a high-competence origin. In total, there are 173 high-high pairs, 78 low-low pairs, and 192 high-low pairs.

In Fig. 5, we plot histograms of the distribution of distances between nearest-neighbouring origins for each of the three categories listed above, with the threshold value chosen to be *p*_0_ = 0.6 (that is, origins are considered “high” if their competence is at least 60%). Fig. 5 confirms that the spread in the distribution of distances for the high-high origin pairs is much greater than for low-low cases. It is clear from the figure that we are much more likely to see large inter-origin separations for the high-high case. In fact, comparing Figs. 5a and 5b, we see that only a very small fraction of pairs have distances greater than 30 kilobases in the low-low case, whereas pairs with distances greater than 40 kilobases are quite common for high-high pairs.

**FIG. 5.**
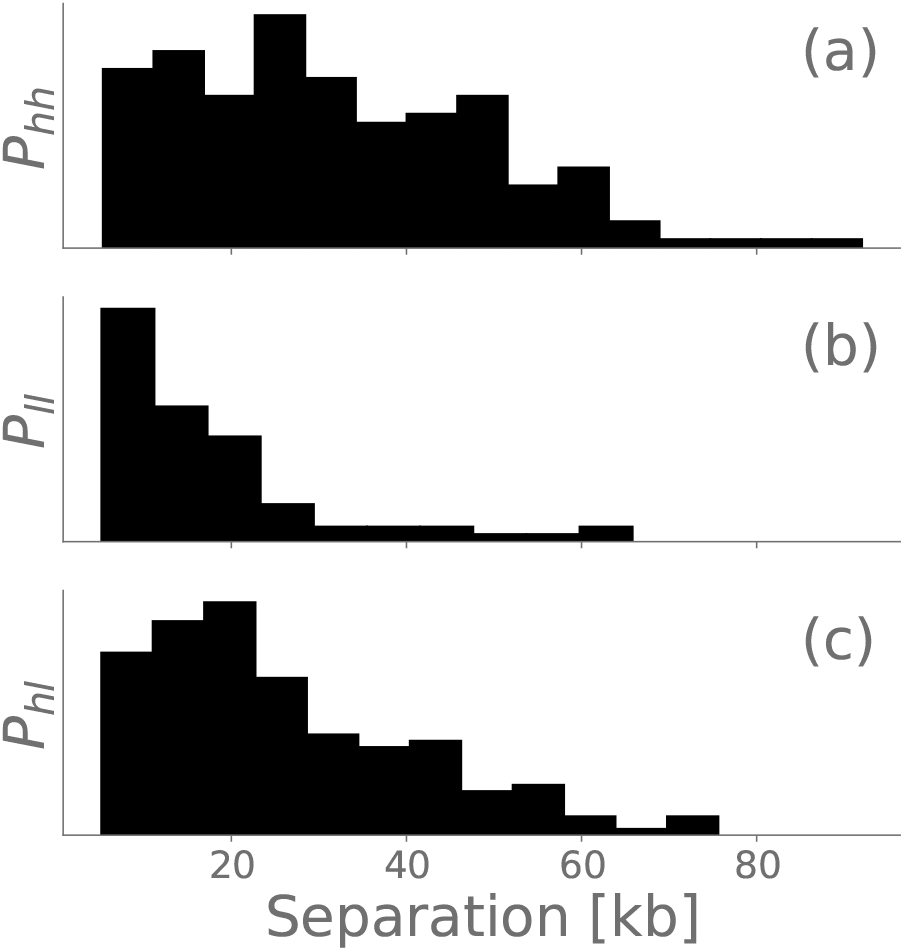
Histograms of the distance between pairs of neighbouring origins, according to their competence. (a) shows data for origin pairs where both origins have high competence (high-high pairs); (b) plots the case where both origins have low competence (low-low pairs); and (c) is the case where one origin has high competence and the other has low competence. For the purposes of this plot, origins with competence greater than 0.6 are considered to have high competence. The values of the frequencies in the vertical axes are not relevant, and are therefore omitted. The origin competence data was taken from [20].

The details of Fig. 5 of course depend on the precise value of *p*_0_, but for any choice of *p*_0_ greater than 0.5, we see a marked difference between the distance distributions for the high-high compared to the low-low pairs, always with the high-high case favouring greater distances between origins.

One may raise the question of whether the correlations between the origin parameters might be a numerical artifact from the fitting process. In order to investigate this, we have simulated replication profiles of populations of artificial virtual chromosomes, with origins whose parameters and positions were all chosen randomly, and then applied the same fitting procedure used to estimate the origin parameters in [20]. The details are in the Appendix. The conclusion is that the fitting of next-generation sequence data does **not** create spurious correlations between origin parameters and their positions that are strong enough to affect our conclusions. We can thus be confident that the correlations shown in 4 and 5 are real.

Figures 4 and 5 provide strong evidence that origin competences and inter-origin distances are correlated in the way predicted by our theory. This is confirmed by calculating the corresponding p-value, which turns out to be 6 *×* 10^*−*13^. This indicates that the correlation is real, and not a product of blind luck. We also use a more sophisticated Bayesian approach to test the significance of the correlation. This is done in the Appendix. The result again confirms that the correlations are real.

We conclude that there is enough evidence of a strong correlation between origin competences and inter-origin distance, and that this correlation is as predicted by our theory.

### Neighbouring origins with higher differences in firing times are more likely to be close to each other

Our theory also predicts correlations between origin activation times and their distances to each other. In Fig. 4b, we plot the inter-origin distance versus the absolute value of the difference in firing times for each nearest-neighbouring origin pair in *S. cerevisiae*, with data taken from [20].

It is a clear from Fig. 4b that neighbouring origins with greater differences in origin firing time tend to be closer to each other, as we predict. The p-value of this data is 10^*−*8^, confirming that the correlation is not a product of chance. We again apply Bayesian inference to assess more rigorously how strongly the origin data supports our hypothesis. The conclusion is again that the correlation is real; for details, see the Appendix.

There is some evidence that the competences and the firing times are correlated [20]: more competent origins tend to fire earlier. This can be seen in Fig. 4c. It is therefore a legitimate question whether the trent in firing time differences we see in Fig. 4b is simply a consequence of the correlation between potentials and inter-origin distances. In order to see if there is statistically significant evidence that the two trends are real, independently of this correlation, we fit a multiple-variable linear regression model, with the inter-origin distances *d* as the response variable, and the mean competences *p*_*av*_ and firing time differences *τ* as the predictor variables. In other words, we fit the statistical model below to the origin pair data:

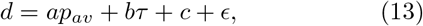

where *c* is the constant term in the regression, and *ϵ* represents noise. The results of this fitting are displayed on the table shown in Fig. 4d. The fact that the p-value for both predictor variables is very low shows that there is statistically significant evidence that both trends are real, and neither can be explained away by the correlation between the two variables.

We have also analysed yeast origin data obtained from the replication model proposed by the Rhind and Bechhoefer groups [23, 31]. Their model does not consider origin competence; they assume that all origins have been licensed at the start of S-phase. Another difference between their model and the one in [20] is that they consider a different set of origins in their model, which has large overlap with the one we use, but is not identical to it. They infer the origins’ average firing times for their model by fitting the model’s predictions to measured replication profiles, and these results are plotted in Fig. 6. The figure shows a negative correlation between the origins’ separations and their firing time differences, in accordance with our prediction. The correlation is not as strong as the one we get from the data in [20], but the p-value is still less than 1%.

**FIG. 6.**
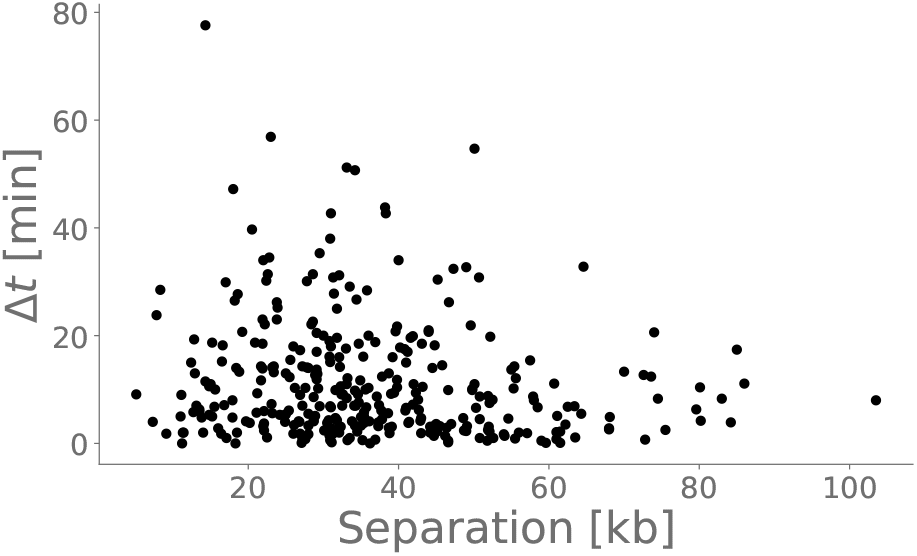
Each point represents one pair of neighbouring origins in *S. cerevisiae*. The horizontal axis is the distance between the origins in both plots. The vertical axis is the difference in average activation time between the two origins. The data was taken from [23].

**FIG. 7.**
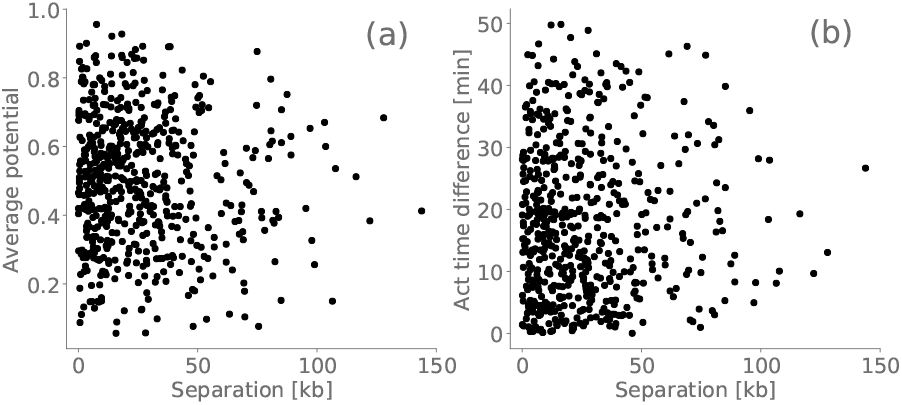
Each point represents one pair of neighbouring origins in simulated chromosomes with randomly generated origins. The horizontal axis is the distance between the origins in both plots. In (a), the vertical axis is the average of the competence of the two origins. In (b), the vertical axis is the difference in average activation time between the two origins.

We conclude that the origin data from both data sets support our prediction that neighbouring origins with higher differences in firing times are more likely to be close to each other than pairs with similar firing times.

## IV. DISCUSSION

Our results support the hypothesis that natural selection favours placements of replication origins in *S. cerevisiae* leading to short replication times. This selection pressure is probably due to the greater vulnerability of cells during DNA replication [24, 25]. Even if the differences in replication time between two competing strains are small, over many generations this can result in one strain thriving and the other going extinct: natural selection is an efficient amplifier of small differences [32].

We note that replication time also plays a crucial role in the *random-gap problem*, which has been extensively investigated in organisms where the locations of origins are not fixed, such as the model organism *Xenopus laevis* [33–35].

A tacit assumption in this work is that the origin locations are a flexible parameter for natural selection to act on, and that mutations resulting in changes in their genome locations are common. This is supported by the fact that events such as genome transpositions are very common in the evolutionary history of most Eukaryotes, including yeast, and they usually lead to origin displacements [26, 36].

If origin locations are selected for their short replication times, why does their distribution not follow the predictions of the theory more closely? The trends seen in Fig. 4 are in the direction predicted by our theory, but the sharp transition from clustered to isolated configurations, predicted by the two-origin model, is nowhere to be seen: although origins with high competence are clearly much more likely to be isolated from nearby origins, some of them do have close neighbours. One of the reasons is the simplicity of our replication model. Real chromosomes and origins are much more complex than our idealised versions of them; to cite just one example, yeast chromosomes typically have dozens of origins, not just two. In addition, there are restrictions in the origin positions which we did not account for in our model; for example, origins usually are not found within genes. And finally, another reason for the imperfect match between our theory and the data is the fact that there are myriad selection pressures acting on the genome which also affect the locations of origins, and are completely independent of the replication timing. So when genes are inserted, deleted or duplicated in the normal course of evolution, distances between origins often change, and sometimes new origins are introduced or existing ones are deleted. The end result is that the overall bias predicted by the theory is still visible, but the fine details predicted by the theory — such as the sharp clustered-isolated transition —are lost.

This study focused on *S. cerevisiae* as a model organism, because it is an organism for which the locations, competences and firing times of all replication origins have been determined. However, many unicellular Eukaryotes have replication origins with fixed positions, and we expect our predictions to hold for these organisms. In addition, many aspects of the replication dynamics are conserved across species, including the locations of most high-competence origins [36].

This paper has focused on the minimisation of replication time as an important selection factor driving the distribution of replication origins. There are other important selection pressures acting on origin positions, however. In [37, 38], the authors focus on a different selection criterion, namely the minimisation of the probability that a double fork stalling event prevents a region of the genome from finishing replication [2]. The resulting “best” origin distribution has the origins regularly spaced from one another, like beads on a string. This may seem at odds with our predictions, since we predict that origins will sometimes bunch together and sometimes stay apart, depending on their competences and firing times; but in fact, the prediction in [37] is what we would expect from our model if all origins had high competence. But the majority of origins in yeast have high competences − 66% of the origins have competence greater than 0.5. The results reported on [37] are therefore broadly consistent with our model, because the high-competence origins dominate the replication dynamics of yeast. This implies that both selection pressures on origin placement — the minimisation of replication time and the minimisation of double fork stalling events — result in broadly similar origin distributions.

As a final remark, we add that our model focuses on the role played by the spatial distribution of origins on the replication time. There are of course other factors that play a role in the evolution of the DNA replication dynamics. For example, the number and density of origins may also change over evolutionary times, and they may also be under selection pressure. But our result is still relevant even then, because it is still true that for a given origin density, some origin distributions lead to faster replication than others. In other words, we are not claiming that our model tries to take into account every single factor that affects the evolution of replication origins; we focus on one important aspect involved in evolution — namely, the positioning of replication origins.

## V. CONCLUSION

We have shown that the positions of replication origins in yeast are strongly correlated with the parameters determining the dynamics of origin firing, namely the competence and average firing time. Specifically, we found that neighbouring origins with low competence are more likely to be clustered together in the genome; and that neighbouring origins with large differences in their firing times are more likely to be clustered. We showed how these correlations are predicted from a mathematical model incorporating the assumption that the positions of replication origins are biased towards configurations minimising the total replication time. This lends support to our hypothesis that the minimisation of replication time is a selection pressure that plays an important role in shaping the spatial distribution of replication origins in the genome of yeast, and potentially in the genomes of other Eukaryotes as well.

## VI. ACKNOWLEDGEMENTS

We want to acknowledge Conrad Nieduszynski for valuable discussions.

For the purpose of open access, the author has applied a Creative Commons Attribution (CC BY) licence to any Author Accepted Manuscript version arising from this submission.

## Appendix A: Quantifying origin parameter fitting bias

Our data analysis was largely reliant on the origin data obtained by fitting data from next-generation sequencing experiments, as described in [19, 20] and [23]. This raises the question of whether the correlations between the origin parameters might be a numerical artifact from the fitting process. In order to eliminate this possibility, we have simulated replication profiles of populations of artificial virtual chromosomes, with origins whose parameters and positions were all chosen randomly. We then applied a global fitting procedure, analogous to the one used to get the origin data in [20]. By randomizing all origin parameters, we ensure that there is no correlation in this “control sample”. If the fitting procedure generated spurious correlations, they would be visible in the result of this fitting. Thus, this is a numerical control “experiment”, to make sure we are not basing our analysis on numerical artifacts.

We generate 20 “in silico” chromosomes with randomly positioned origines, whose parameters mimick the ones found in yeast: the mean and standard deviations of the firing times are taken from a normal distribution with the same statistical properties of those found in yeast. The potentials of each origin was chosen randomly with a uniform distribution between 0 and 1. The fork velocity was fixed at 1.6 kb/min. The snapshot times for the replication profiles (see [20] for details) were chosen as t=10,30,50,70, and 90 minutes. Each chromosome in the virtual population is also chosen with a random length with the same average and standard deviation as in yeast, and the number of origins for each chromosome is chosen to that the origin density is the same as that of yeast. The same fitting procedure is applied as in [20], and the results are plotted in Figs. S2(a) and S2(b).

Noo correlations can be discerned in these figures — contrast them with Figure 3 in the paper. The Pearson correlation coefficient is 0.02 for the activation time differences versus distance data, with a p-value of 0.55, indicating no statistically significant correlation. This is in contrast to the correlation of -0.29 and p-value of 10^*−*8^ found in the data.

For the average potential versus distance data, the correlation is a bit higher, with a value of -0.08, and a p-value of 4%, possibly indicating a very weak bias — although the p-value is marginal. If the slight bias is real, it definitely does not explain away the very strong correlation found in the data — a coefficient of 0.33 and a p-value of 6 *×* 10^*−*13^.

We conclude that the fitting of next-generation sequence data does **not** create significant spurious correlations between origin parameters and their positions, and that the correlations we found are real, and not mere artifacts of the fitting procedure.

## Appendix B: Bayesian inference applied to the origin data

### 1. Bayesian model for the correlation between distance and competence

In order to test the hypothesis that low-competence origins tend to cluster together, we first formulate a very simple statistical model for the correlation between the distance between origin pairs and their competences. First we separate the origin pairs into “high-competence”, “low-competence”, and “mixed” pairs. An origin pair *i* is a high-competence pair if both origins have competence greater than a threshold value *p*_0_; *i* is low-competence if both origins have competence less than *p*_0_; and it is mixed if it is neither high-competence nor low-competence. We set the threshold competence *p*_0_ to 0.6 for the purposes of this analysis; although this choice is somewhat arbitrary, it is close to the transition probability predicted by the mathematical model presented above. The results of our analysis is largely independent of the precise value assigned to *p*_0_.

Our first statistical model assumes that the neighbouring origin distances *s*_*i*_ are described by an exponential distribution:

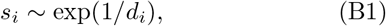

where the mean distance *d*_*i*_ depends on whether the origin pair is high-competence, low-competence or mixed:

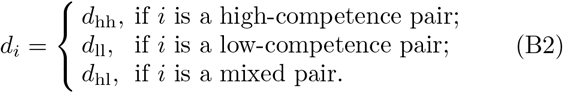

This model therefore describes the origin distances as a mixture of three exponential distributions. *d*_hh_, *d*_ll_ and *d*_hl_ are the three parameters of the model. The prior distribution consists of a uniform distribution of *d*_hh_, *d*_ll_ and *d*_hl_ over the interval [0, 100] kilobases.

Applying Bayesian inference using Monte-Carlo methods [39], we can infer from the data the probability distributions (that is, *posterior* distributions) for the distance parameters *d*_hh_, *d*_ll_ and *d*_hl_ from the origin data.

The results of this analysis for *d*_hh_ and *d*_ll_ are shown in Fig. 8a. If the competences and inter-origin distances were not correlated, we would expect the two histograms to be identical (other than noise due to the finite sampling size). Instead, we can see that the two distributions are clearly separated, with almost no overlapping. This again strongly supports the correlation our model predicts. The probability that the data is the result of pure chance, and that no correlation exists, can be estimated from the area of the overlap between the two distributions in Fig. 8a, and is approximately 10^*−*4^.

**FIG. 8.**
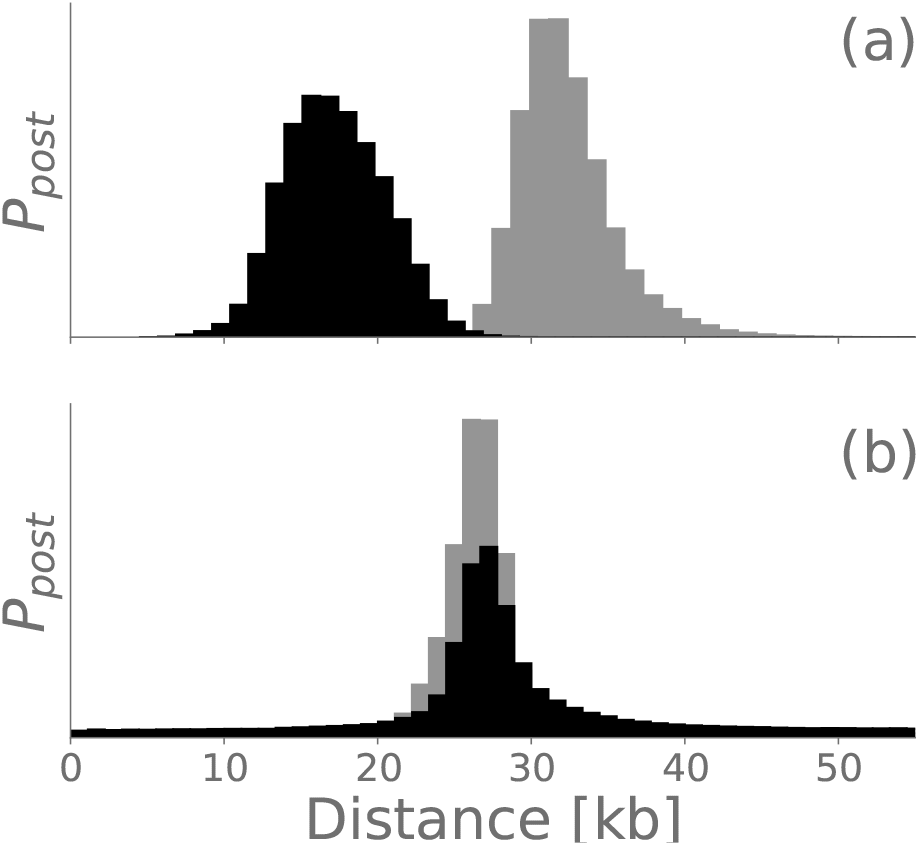
Posterior probability distributions for the parameters *d*_hh_ (black) and *d*_ll_ (grey) in Eq. (B2), obtained by running the Hamiltonian Monte Carlo method described in the Methods section. (a) shows the results for the *S. cerevisiae* origin data; (b) shows the result of a control data set, where the origins are subject to a random permutation, destroying any correlations present in the actual data. The values of the probability densities in the vertical axes are not relevant, and are therefore omitted.

For comparison, Fig. 8b shows the same analysis done on a modified version of the origin data where the origins were kept at the same positions, but their competences were shuffled and reassigned to all origins at random. This has the effect of destroying any existing correlation between origin positions and their competences. As expected, Fig. 8b shows that the inferred posterior distributions of *d*_hh_ and *d*_ll_ completely overlap, showing that in the shuffled data, the inter-origin distances are not affected by their competence. This is in sharp contrast to the analysis of the real data shown in Fig. 8a.

### 2. Generalised model

In the model described above, we had to postulate a somewhat arbitrary partition of origins between high- and low-competence. So we also show the results using a different statistical model, which does not rely on this partition. This model is also more general, because in addition to the distance-competence correlation, it also takes into account the influence of the firing time difference.

The model again assumes an exponential distribution for the inter-origin distances, with a linear dependence of the exponent on the average competence and firing time difference of the pairs:

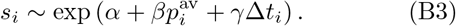

The correlations are now encoded in the parameters *β* and *γ*. The priors for the three parameters *α, β* and *γ* are uniform distributions over the interval [*−*20, 20].

This alternative Bayesian analysis does not require choosing an arbitrary threshold value *p*_0_, and has the added advantage of unifying the analysis of the competence and the firing-time dependencies. The parameters *β* and *γ* describe how correlated the average competence and the firing time difference are to the inter-origin distance, respectively. Positive values indicate anti-correlation (that is, the distances grow with the quantity), whereas negative values indicate positive correlation (distances decrease as the quantity grows); values around zero indicate the absence of correlations.

Using Bayesian inference, we can find the posterior distributions of this model’s parameters from the origin data. Figure 9b shows the resulting distribution for the *β* parameter. The histogram lies almost completely on the negative side, indicating *positive* correlation — that is the inter-origin distance increases with the origin competence, as predicted.

**FIG. 9.**
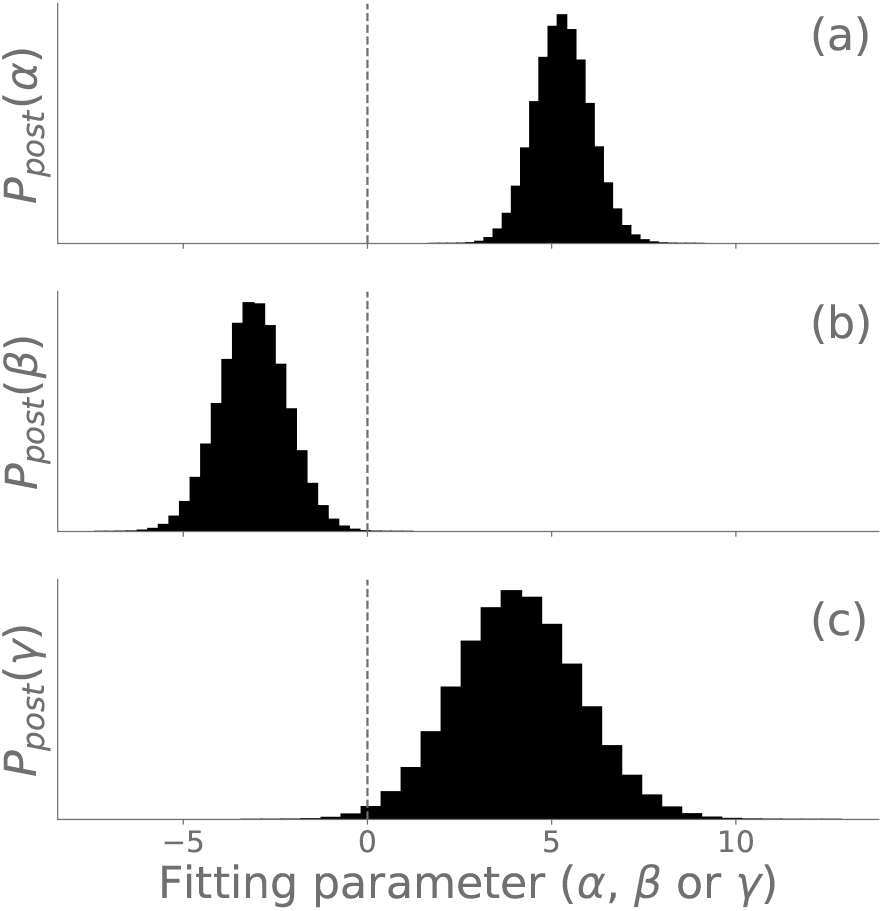
Posterior probability distributions for the parameters *α* (a), *β* (b) and *γ* (c) in Eq. (B3), obtained by running the Hamiltonian Monte Carlo method described in the Methods section. The values of the probability densities in the vertical axes are not relevant, and are therefore omitted.

Figure 9c shows the result of Bayesian inference applied to the linear model for the parameter *γ*, which describes the influence of the firing time differences on the inter-origin distances. We can see that, although a small portion of the left tail of the histogram lies on the negative side of the axis, most of the distribution lies on positive values of *γ*. This shows a *negative* correlation between origin distances and firing time differences, just as predicted by our theory.

### 3. Monte Carlo simulation of the Bayesian models

We use the Hamiltonian Monte Carlo (HMC) method [39] to sample the posterior distribution of the statistical models described above. The HMC method converges very quickly to the posterior distribution, and is ideal for our purposes. We use the Stan language [39, 40] through its Python API [41] to do all the Bayesian analysis. In the simulation of both statistical models described above, we used 8 chains with 2 *×* 10^5^ iterations per chain, excluding the warmup phase of 2 *×* 10^5^ iterations. The effective sample size was found to be over 10^5^ in every case. Convergence was monitored through scale reduction parameter 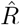, which we ensured was equal to the convergence value of 1 to within at least three decimal places in every run.

## Notes

### Competing Interest Statement

The authors have declared no competing interest.

### Summary of Updates

Version submitted to PRE. The only change compared to the previous version is that the supplementary material has been merged into the body of the paper.

